# Positive selection and structural disruption of KCNA5 by mutation in colorectal cancer

**DOI:** 10.64898/2026.10.09.758060

**Authors:** Binglin Xu, David Shorthouse

## Abstract

Ion channels regulate cell proliferation, survival and migration, but the functional consequences of many ion channel mutations in cancer are unknown, and electrophysiological characterisation at scale is impractical. Here we use the mutational landscape of colorectal adenocarcinoma as a functional filter to identify potassium channel mutations under selection. A dN/dS screen of 16 potassium channels expressed in the large intestine identified KCNA5 (Kv1.5) as the strongest candidate under positive mutational selection. These mutations cluster in the T1 tetramerisation, S1–S4 voltage-sensor and pore domains, yet avoid the selectivity filter, indicating that selection does not favour loss of conduction. Within the pore region, the observed mutations occupy a biophysical space distinct from known C-type inactivation, activation-gate and toxin-binding mutations. In the voltage sensor, mutations affect gating-charge arginines. In the T1 domain, mutations fall on inter-subunit interfaces conserved across the KCNA family, and saturation mutagenesis energy calculations predict that they destabilise subunit contacts or the folding of the domain. Molecular dynamics simulations of D166N and G128R show loss of coordinated interface dynamics and distortion of the local backbone, respectively. Transcriptomic analysis of KCNA5-mutant tumours shows reduced expression of cell-cycle, KRAS-related and inflammatory pathways, and we observe differences between transcriptional effects induced by T1 and S1-S4 domain mutations. These changes are not reproduced by low KCNA5 expression alone, consistent with an effect specific to the mutant protein. Together, these computational results highlight that KCNA5 is under mutational selection in colorectal adenocarcinoma, and that these mutations are likely functional and induce phenotypic effects on tumors.

## Introduction

Ion channels are pore-forming membrane proteins that allow selective ion movement across the lipid bilayer. The resulting changes in membrane potential and electrical activity trigger signalling cascades that regulate core processes in cancer, including cell-cycle progression, evasion of apoptosis, and migration and invasion, and the link between ion channels and cancer has attracted considerable interest[1,2]. Potassium channels have particularly diverse relationships with cancer and have been reported to drive or repress oncogenic processes in several tumour types [3,4]. Examples include Kv11.1 (hERG1), which interacts with integrin signalling[5], and KCNQ channels, which participate in Wnt signalling to regulate proliferation, differentiation and migration [6].

Potassium channels are central to the physiology of the intestinal epithelium, where they regulate membrane potential and drive potassium secretion – helping to maintain epithelial polarisation[7], and several have established roles in colorectal cancer. KCNQ1 has been shown to be prognostic in colon cancer - where its loss predicts disease recurrence[8], it interacts bidirectionally with β-catenin in CRC cell differentiation[9], and cancer-associated loss-of-function mutations in KCNQ1 enhance Wnt/β-catenin signalling[10]. Related work from our group also shows that KCNQ channels modulate Wnt activity in gastro-oesophageal adenocarcinoma[6]. KCNQ1 illustrates both that potassium channel mutations can be functionally relevant in gastrointestinal cancer and that their effects are mediated through core oncogenic pathways. Given the large size and diversity of the potassium channel family, we sought to study other members that may be acting in similar ways.

Among candidate channels, KCNA5 (Kv1.5) is of particular interest. It is a Shaker-family voltage-gated channel that carries the ultra-rapid delayed rectifier current in human atrial myocytes[11], where loss-of-function mutations cause atrial fibrillation[12]. A role in cancer has been previously studied: KCNA5 silencing by promoter methylation supports proliferation in Ewing sarcoma[13], and Kv1.3 and Kv1.5 more generally have been linked to cell-cycle progression in a number of cancers[14,15]. This is consistent with the long-standing link between K⁺ channel-driven membrane hyperpolarisation and progression through G1/S stages of the cell cycle[16,17]. Like other Kv1 channels, KCNA5 is a homotetramer, each subunit comprising a cytoplasmic T1 tetramerisation domain, a voltage-sensing domain (S1-S4) and a pore domain (S5-S6) containing the selectivity filter. The T1 domain governs subunit assembly and subfamily-specific recognition[18], and its polar inter-subunit interface is coupled to conformational changes that open the channel (D. Minor & Berger, 2000).

Studying these consequences in cancer is challenging. Electrophysiology is technically demanding, and the many channel subtypes expressed together in a membrane make it hard to attribute an effect to any single channel. Activation mechanisms vary across subtypes (voltage, ligand or mechanical force), which prevents a unified experimental approach, and effects are cell-type dependent, so that the same channel can promote proliferation in one tissue and apoptosis in another[20]. Somatic mutations in ion channel genes are common across cancers, but their consequences are mostly uncharacterised[21], and functional characterisation of individual cancer-associated channel variants remains rare[22].

Tumour genomes offer a complementary route. Where cancers evolve under positive selection, mutations that arise more often than expected under neutral evolution, after accounting for mutational context[23,24], are unlikely all to be passengers. Structure-based analysis of missense mutations has been used to identify functionally important regions in many cancer gene[25,26], including by our group [27–29]. Tumours thus act as a “functional filter” for mutations, and the pattern of selected mutations is a readout of protein function that does not depend on real-time electrophysiology. Combining selection analysis with structure-based analysis and molecular dynamics allows testable mechanistic hypotheses to be generated at residue level. This is particularly useful for voltage-gated potassium channels, for which the structural and functional framework is well characterised.

Here, we screened potassium channels expressed in the large intestine for positive mutational selection in colorectal adenocarcinoma and identified KCNA5 as the strongest signal. We show that its missense mutations cluster in the T1, voltage-sensor and pore domains but avoid the selectivity filter, and that mutations in the pore region occupy a distinct biophysical space from known gating-altering mutations. We use saturation mutagenesis and molecular dynamics to predict that T1-domain mutations destabilise inter-subunit contacts and alter the dynamics of the interface. Finally, we examine the transcriptional profile of KCNA5-mutant tumours and compare mutations in different domains.

## Results

### KCNA5 is under positive mutational selection in colorectal adenocarcinoma

To identify potassium channels under mutational selection in colorectal cancer, we applied dNdScv[23] to colorectal adenocarcinoma samples in COSMIC (v103)[30], restricting the analysis to 16 potassium channel genes expressed in the large intestine[31]. dN/dS compares the observed ratio of nonsynonymous to synonymous mutations with the ratio expected under neutral evolution, correcting for trinucleotide context and gene-specific mutation rate. Values above 1 (log2 > 0) indicate an excess of nonsynonymous mutations consistent with positive selection, and values below 1 a deficit. KCNA5 showed the strongest evidence of positive selection for missense mutations (p = 0.0001, **Fig 1A**). KCNK12 showed the strongest evidence of negative selection (fewer mutations than expected by chance, p = 0.0005), we found no truncating mutations in KCNA5 in our cohort – so selection acts on missense mutations rather than just on loss of the gene product. We therefore focussed our analysis on KCNA5. Overall in The Cancer Genome Atlas (TCGA)[32] – which makes up a portion of the COSMIC database, with matched copy number data, KCNA5 is mutated in 6% and amplified in 1.7% of 583 colorectal adenocarcinomas.

**Figure 1:**
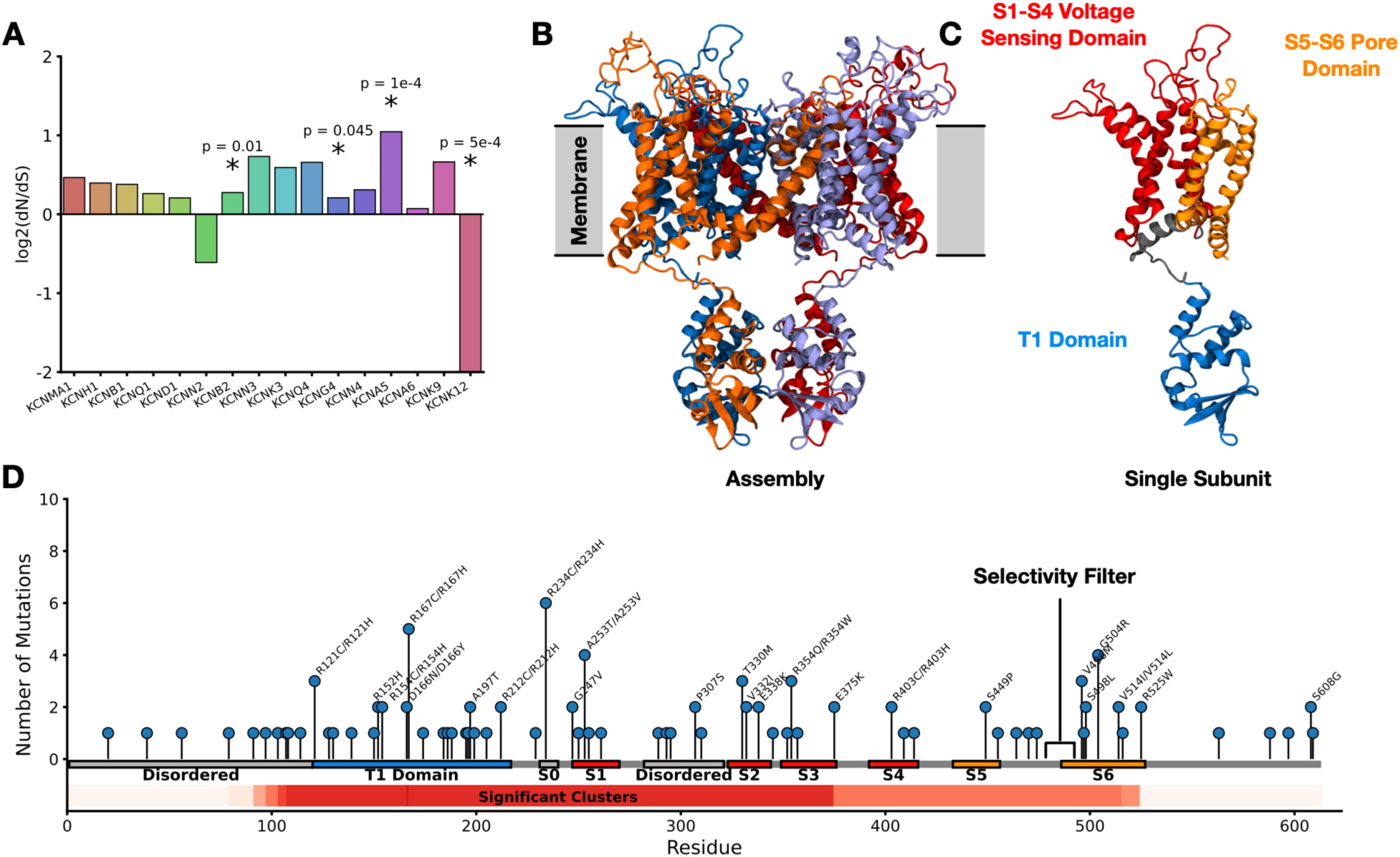
KCNA5 is under mutational selection in colorectal adenocarcinoma. A) dN/dS analysis of potassium channels expressed in the colorectal tract * represents Bonferroni corrected p-value <= 0.05. B) Structure of the full assembly of KCNA5 with each subunit coloured differently, C) Structure of a single subunit of KCNA5 highlighting the different functional domains. D) Lollipop plot of mutations observed in KCNA5 in the COSMIC cohort, with NMC clustering analysis results included (below).

The 103 missense mutations in the COSMIC cohort are expected to be enriched, but not composed entirely, of functionally relevant mutations: dNdScv estimates that 52.4% of them are due to selection. Because mutations selected in cancer are likely to influence channel activity, their locations give a view into how KCNA5 contributes to tumour fitness and into how mutations alter protein function.

We next examined the distribution of these mutations along the protein. Non-random mutational clustering (NMC)[33], with the permutation background restricted to the structured region (residues 119–527), identified hundreds of 1D statistical clusters at corrected p ≤ 0.05 (**Fig 1D**). They localise to the T1 tetramerisation domain, the S1–S4 voltage-sensor domain and the S5–S6 pore domain. Most mutations in the voltage-sensor and pore domains lie on helices.

### Colorectal cancer mutations cluster around the selectivity filter and are biophysically distinct from C-type and Activation gate mutations

Despite the concentration of mutations observed in and around the pore domain, no mutations were observed within 5.5 Å of the selectivity filter (**Fig. 2A, B**). We tested the significance of this absence using two Monte Carlo simulations of random mutagenesis (**Figure S1**): permuting all 103 mutations across the full 613-residue sequence yielded p = 0.0213 for likelihood of observing no mutations in this region, while restricting the permutation to the 77 positions between the first and last pore-domain mutation - giving a higher local mutation density of 21 mutations, yielded p = 0.0001. The absence of selectivity filter mutations is therefore unlikely to due to random chance. Instead, this analysis suggests purifying selection acts to remove mutations that would abolish ion conduction, suggesting that the selective advantage conferred by the observed mutations is likely not purely due to a blocking or abolishment of channel function.

**Figure 2:**
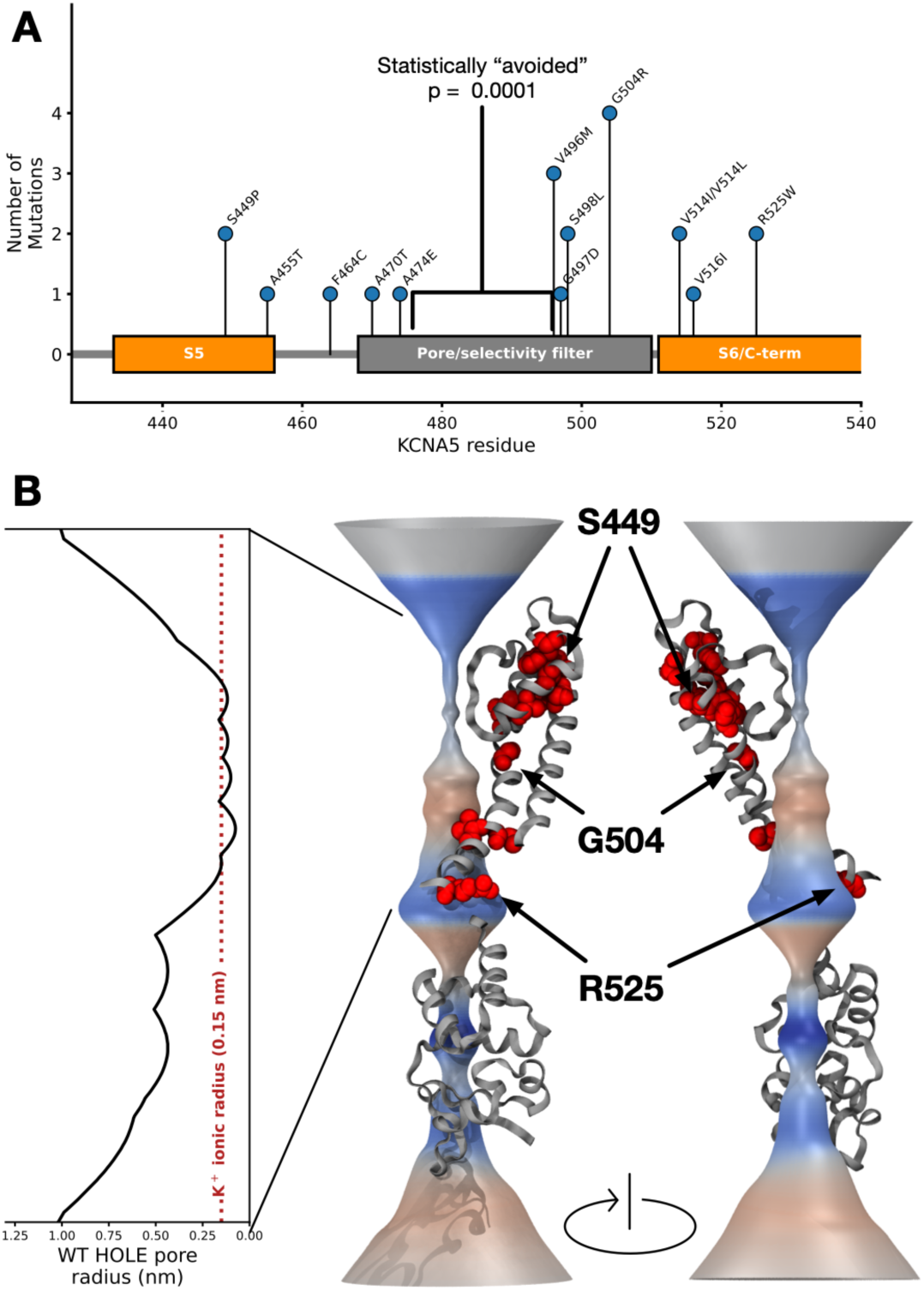
Mutations in KCNA5 in colorectal adenocarcinoma statistically avoid the selectivity filter. A) Lollipop plot of KCNA5 with observed mutations, statistically avoided region around the selectivity filter is highlighted. B) Render of the pore profile of KCNA5 calculated with HOLE, with mutations observed in colorectal cancer highlighted in red.

Mutations to the pore and selectivity filter of potassium channels can impact gating in a number of ways – they may completely ablate the channel activity by damaging the selectivity filter structure, but they can also alter the kinetics and voltage-dependence of C-type (slow) inactivation without eliminating conduction altogether. Residues immediately flanking the filter such as Thr449 in Shaker and its Kv1 equivalents (KCNA5 equiv. Arg487) are highly sensitive to substitution. Charge-introducing mutations at this location accelerate inactivation up to 100-fold, while its removal can abolish inactivation entirely[34]. Turret-loop substitutions produce similar effects, whereby histidine mutations in KCNA5 (His463) accelerate inactivation regardless of the charge introduced[35] - consistent with the hydrogen-bonding network linking the pore helix to the turret that acts as a molecular timer for filter collapse[36].

A second, mechanistically distinct class instead alters ion selectivity largely independent of gating kinetics. Pore-helix substitutions beneath the filter, such as V370G in Kv1.2 (KCNA5 Val476), collapse K+/Na+ discrimination, producing large depolarizing shifts in reversal potential and substantial non-K+ flux[37]. Mutagenesis of the signature sequence itself confirms selectivity and inactivation can be uncoupled as separate phenomena[38,39].

To better characterise these mutations, we calculated a biophysical fingerprint for every possible mutation in the selectivity filter/pore region of the protein (defined as between residues 433 and 527) (**Fig 3A**). For every possible mutation in this region we calculated the energetic changes induced by mutation (through a saturation mutagenesis screen performed using Foldx), changes in the pore profile as calculated by HOLE[40] and CHAP[41], distances of key hydrogen bonding interactions known to drive gating activity in mutation models, and elastic network model properties describing each position’s intrinsic flexibility and dynamic coupling to the selectivity filter in the wild-type structure. This complex fingerprint comprising 21 features was then analysed using Principal Component Analysis (PCA). The top two Principal Components explained only 23.4% and 10.7% of the total variance, respectively (34.1% combined), indicating that no single dominant axis captures the biophysical impact of mutation in this region and that a substantial fraction of the variance is distributed across many weakly-correlated properties. Despite this, overlaying the mutations observed in colorectal adenocarcinoma, as well as a further literature-derived set of mutations involved in other gating alterations shows a separation of between the cancer observed mutations and other gating altering mutations. Statistical analysis shows a significantly large separation of the 24 Principal Component centroids of the cancer observed mutations to all other groups of mutations (Cancer vs C-type inactivation: p = 0.0002, Cancer vs Toxin binding altering: p = 0.0032, Cancer vs Activation gate p = 0.0078), and was statistically unlikely (p = 0.020) compared to the mutational background.

**Figure 3:**
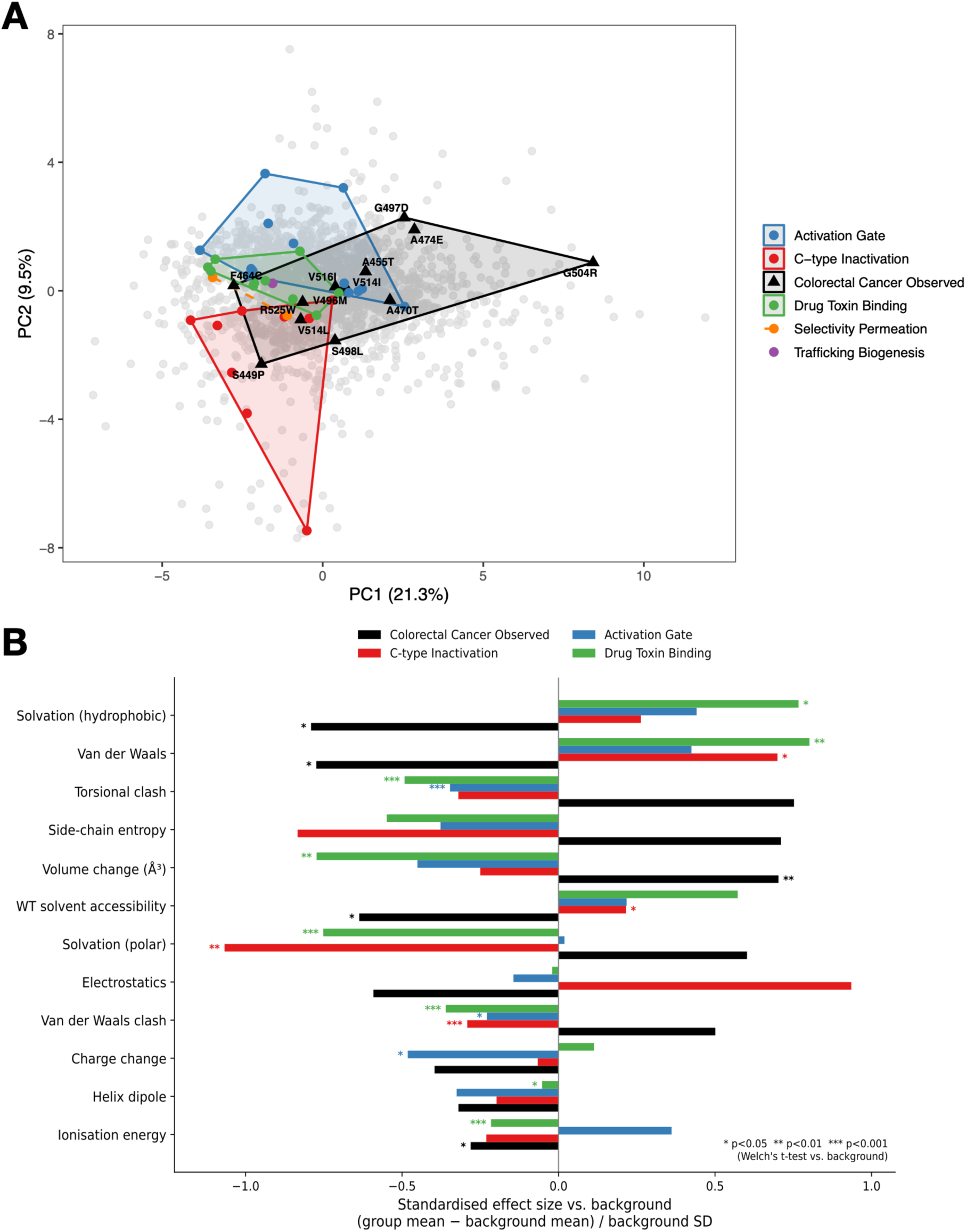
KCNA5 selectivity filter/gating region mutations occupy a novel biophysical mutational class. A) Principal Component Analysis of the full mutational landscape with mutations observed in Colorectal Cancer (black), and other biophysical classes annotated. Mutational regions are highlighted with convex hull analysis. B) Feature extraction of the differences in biophysical characteristics for the 4 major mutational classes against the saturation mutation background. * p <0.05, ** p<0.01, *** p<0.001. P represents Welchs t-test.

Feature contributions show cancer mutations in this region occupy a systematically distinct region of biophysical property space relative to mutations characterised by electrophysiological and pharmacological studies from the literature (**Fig 3B**), skewing toward larger, more sterically disruptive substitutions (increased side-chain volume, torsional strain, and van der Waals clash) rather than the solvation- and packing-driven signatures characteristic of known C-type inactivation, activation-gating, or drug-binding mutations. Overall this points to mutations in this region observed in colorectal cancer occupying a distinct biophysical class not previously observed.

### Mutations in the S1-S4 helix of Kv1.5 are likely to influence gating

One of the most characterised areas of all potassium channels are the S4 helices. These function as regulators of channel activity through the paddle-like movement of the helix in the membrane in response to voltage changes. Classic experiments have shown that altering the 6 charge bearing arginines on these helices has large influence on gating activity, and that alterations to the ability of this helix to move has consequences on the gating ability of the channel. Within our cohort we find a large number of mutations within the S1-S4 helical bundle (**Fig 4**), with some landing directly on the charge bearing arginines involved in gating. The most frequently mutated position in this group is A235, which interacts with the S4 helix, and is predicted by FoldX[42] to have a high folding gibbs free energy change – indicating an induction of instability (Calculated DDG for A235T – 2.17kcal/mol).

**Figure 4:**
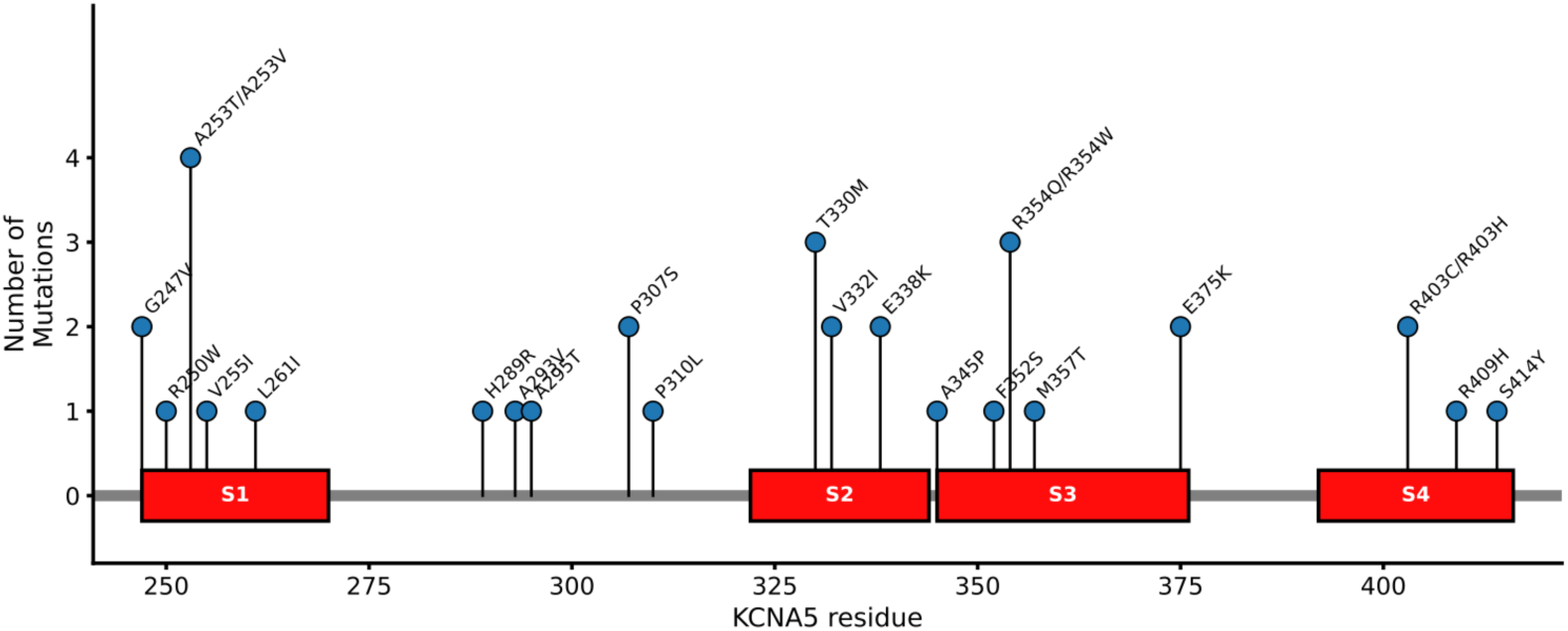
Lollipop plot of the S1-S4 region of KCNA5 with mutations in our cohort overlayed.

Interestingly we observed mutations on two of the gating charges directly R403H/C (R2), and R409H (R4). Loss of these gating charges would be expected to impair voltage-dependent activation of the normal K⁺ conductance[43–45]. The substitution to histidine observed here is of particular interest: in Shaker, the equivalent substitutions do not merely neutralise gating charge but create a distinct, aberrant proton-conducting pathway through the voltage-sensor domain itself (a “gating-pore” or “omega” current), separate from the channel’s canonical K⁺-selective pore[46–48]. This class of gating-pore leak current is an established disease mechanism in other voltage-gated channelopathies (e.g. hypokalemic periodic paralysis via homologous Nav/Cav S4 mutations),[49–51] raising the possibility that R403H and R409H combine loss of normal voltage-sensor function with gain of a novel, aberrant ionic leak - though this has only been demonstrated previously in Shaker.

### Colorectal cancer mutations destabilise inter-subunit contacts in the T1 domain and are associated with altered gating

We next studied mutations in the T1-domain of KCNA5 – where there are a number of recurrent sites and evidence of clustering (**Fig 5A**). Examining the locations of the T1-domain mutations in detail uncovers that some key mutations are located on the subunit interfaces. Each T1 domain presents two interaction surfaces to adjacent subunits, termed the A side and B side [52] (**Fig 5B**); in the tetrameric assembly, the principal contacts are formed by the A3 helix and B2 sheet on the A side, the B3-A3 and B1-B2 loops on the B side, and the A3-A4 loop on both sides. Several of the observed mutations fall directly on these interaction surfaces (**Fig 5C**). In particular, R167, D166, and G128.

**Figure 5:**
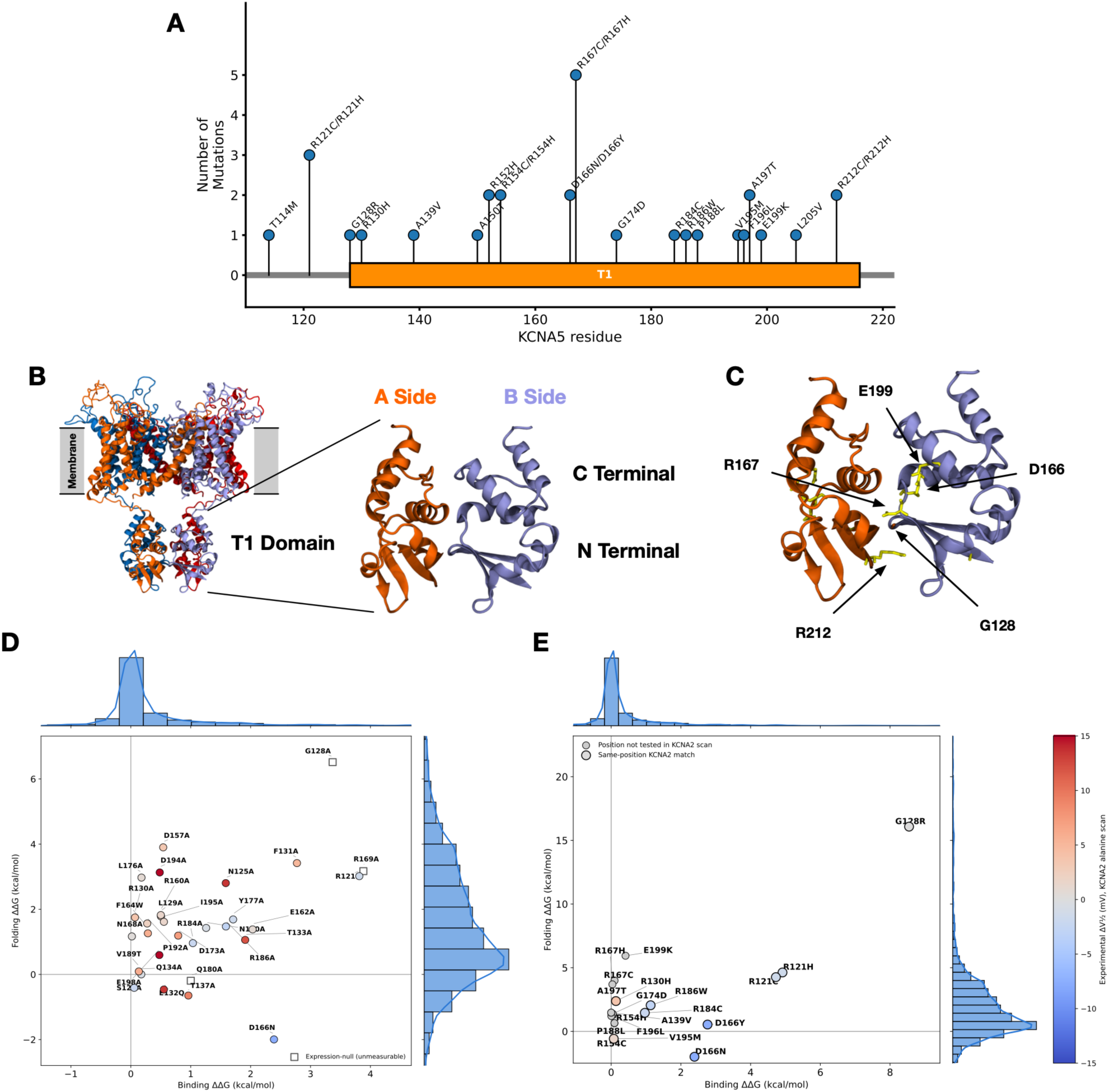
KCNA5 mutations to the T1 domain influence intersubunit binding and folding. A) Lollipop plot of mutations in our cohort in KCNA5. B) Structure of KCNA5 with T1 domain highlighted. C) Structure of T1 domain with recurrently mutated positions highlighted. D) Folding and Binding energy calculations for mutations studied by alanine scan in Minor Jr et al. Points are coloured by their experimentally determined change in activation half life. E) Folding and Binding energy calculations for mutations in our cohort. Points are coloured by their experimentally determined change in activation half life.

Two interface residues in particular - R121 and D166, mutated in 3 and 2 patients respectively, each contact at least three residues across the subunit interface (**Fig 5C**), contacts that are conserved across the KCNA family and have been characterised in the homologous channel KCNA2[19]. Notably, alanine substitution at the equivalent positions in KCNA2 produces a hyperpolarising shift in activation V1/2[52], suggesting that disruption of these contacts alters the energetics of voltage-dependent gating of KCNA5.

To assess the energetic consequences of the observed T1-domain mutations more systematically, we used FoldX to calculate both the change in inter-subunit binding energy (binding ΔΔG) and folding energy (folding ΔΔG) for every possible amino acid substitution in the T1 domain, generating a background distribution of mutational effects against which the cancer-associated mutations could be compared. We also compared our results to the equivalent mutations assayed experimentally by an alanine-scanning mutagenesis screen of the homologous interface in KCNA2 (**Fig 5D**)., for which electrophysiological measurements of the shift in voltage-dependence of activation (ΔV½) are available. Three of the experimentally characterised positions are of particular interest. G128A abolished channel expression entirely (non-functional channel) and was correspondingly predicted to substantially destabilise both the fold (folding ΔΔG = +6.52 kcal/mol) and the inter-subunit interface (binding ΔΔG = +3.37 kcal/mol). D166N produced the largest measured shift in activation voltage of the screen (ΔV½ = −7.72 mV) and was predicted to destabilise the interface (binding ΔΔG = +2.39 kcal/mol), though interestingly it was predicted to be mildly stabilising for the fold itself (folding ΔΔG = −2.00 kcal/mol), suggesting its functional effect may be related to stabilising the fold and altering the dynamics of its host subunit, as well as effects on the interface. R121L also produced a comparatively modest activation shift (ΔV½ = −1.56 mV) and was predicted to destabilise both binding (+3.81 kcal/mol) and folding (+3.02 kcal/mol).

Mapping the mutations observed in our own cohort onto the same binding–folding landscape (**Fig 5E**), we find that G128R is predicted to be the most severely destabilising mutation in the cohort on both axes simultaneously (binding ΔΔG = +8.55 kcal/mol; folding ΔΔG = +16.08 kcal/mol), exceeding its alanine-substituted counterpart at the same position. R121 mutations (R121H, R121C; n=3 patients) show a similar dual-axis destabilisation (binding ΔΔG = +4.73 to +4.92 kcal/mol; folding ΔΔG = +4.26 to +4.63 kcal/mol), consistent in character with the experimentally modest but structurally destabilising R121L. By contrast, R167 - the most recurrently mutated residue in our cohort (R167C, R167H; n=3 patients) shows a distinct, asymmetric signature: negligible predicted effect on interface binding (ΔΔG = +0.04 to +0.10 kcal/mol) alongside substantial predicted fold destabilisation (ΔΔG = +3.69 to +3.99 kcal/mol), suggesting R167 mutations may act specifically through protein misfolding rather than disruption of the T1–T1 inter-subunit interface.

We also examined where the four most recurrently mutated T1 positions in our cohort (R121, G128, D166, R167) sit relative to the hinge sites of the slowest five calculated elastic network (GNM) modes. Hinges in these models being near-zero-fluctuation pivot points around which the surrounding structure moves, positions where a local change in rigidity has disproportionate effects on global motion. While the single slowest mode (mode 1) was dominated by motion outside the T1 domain (hinges at residues 225 and 526/527), two subsequent slow modes (modes 2 and 4) described motions localised to the T1 domain itself. All four cohort positions fell within 1-3 residues of one of these identified hinges (**Fig S2**). This is consistent with a mechanism in which these mutations act by perturbing the local rigidity.

To characterise the structural consequences of T1-domain mutations at atomic resolution, we selected two mutations for MD simulation: D166N, which corresponds to the position producing the largest hyperpolarising shift in ΔV½ in the KCNA2 alanine scan as well as being predicted to stabilise the individual T1 subunit whilst disrupting the interface, and G128R, which produces the largest predicted ΔΔG in both dimensions of the FoldX analysis and renders the homologous KCNA2 channel non-functional. Both of these residues were also identified as being near to hinge-sites in our elastic network modelling. Mutant structures were generated by homology modelling and simulated alongside wild-type KCNA5 in a POPC lipid bilayer.

In the wild-type structure, the carboxylate group of D166 bears two resonance-stabilised oxygens that act as hydrogen bond acceptors, forming contacts with T131, T137, Q134, and Q180 across the subunit interface and stabilising the local interaction network (**Fig 6A**). The D166N substitution removes the negative charge and replaces the carboxylate with a neutral amide, which can participate in hydrogen bonding but offers only one carbonyl acceptor and introduces a donor, resulting in geometrically more restricted interactions.

**Figure 6:**
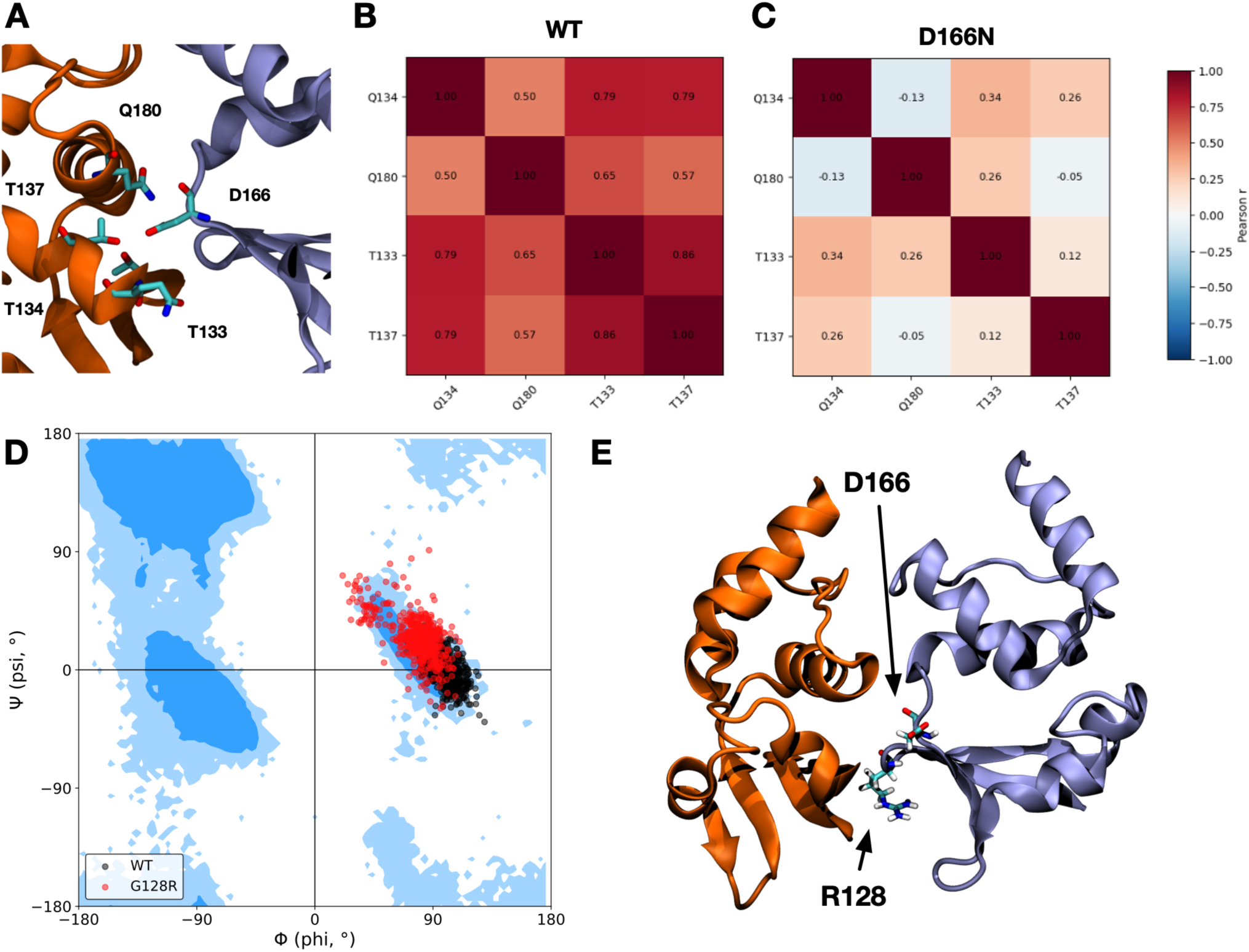
Molecular Dynamics simulations reveal dynamic effects of mutations to the KCNA5 T1 domain in colorectal cancer. A) Position and structure of residue D166 in the T1 domain A/B side interface. B) Co-correlation of molecular movement of residues in the interface for MD simulations of the WT structure. C) Co-correlation of molecular movement of residues in the interface for MD simulations of the D166N structure. D) Ramachandran plot of G128 position in WT and G128R molecular dynamics simulations. E) Screenshot from G128R md simulation showing location of R128 in relation to D166.

MD simulations confirmed this predicted disruption. Inter-residue distances between position 166 and its four interaction partners were predominantly 3.5-4.0 Å in the wild-type, consistent with stable hydrogen bonding, with a secondary population above 4.0 Å reflecting the transient, dynamic character of the native interface. In the D166N mutant, this bimodal distribution collapsed to a unimodal one: T131 and T137 lost contact with N166 entirely, while Q134 and Q180 shifted to alternative interactions - OE1 of Q134 to NE2 of N166, and NE2 of Q180 to the backbone carbonyl of N166 (**Figure SX**). The loss of the longer-distance peak indicates that the mutant interface is conformationally trapped rather than dynamic.

To assess whether this conformational trapping reflects local side-chain rearrangement alone or a broader change in T1 domain dynamics, we computed pairwise Pearson correlations between these inter-residue distances across the simulation. In the wild-type, all pairwise correlations exceeded r = 0.5 (up to r = 0.86; p < 10⁻¹⁵⁰), indicating that interface residues move in a concerted, domain-scale fashion coupling their motion together (**Fig 6B**). In D166N, correlations fell sharply (r = −0.13 to 0.34): with each contact now locked into a single state so residue fluctuations become effectively independent of one another (**Fig 6C**). This shows that the D166N mutation does not simply weaken individual contacts, but disrupts the coordinated dynamics of the T1 interface as a whole, locally rigidifying each contact, consistent with the folding energy calculations suggesting a stabilisation of the T1 domain.

G128 occupies a type II β-turn in a tightly packed region of the T1 interface. The introduction of arginine at this position is predicted to restrict backbone dihedral angles incompatible with the β-turn geometry and to sterically perturb the surrounding β-sheet packing. MD simulation confirmed this: across the full trajectory, the wild-type backbone dihedral angles at residue 128 occupied the canonical β-turn region (Φ = 99.1 ± 9.3°, fi = 0.7 ± 12.2°), a conformation normally accessible only to glycine; in the G128R mutant this shifted significantly (Φ = 78.7 ± 15.3°, fi = 24.0 ± 15.6°), with markedly greater variance, consistent with the bulkier arginine side chain being unable to comfortably occupy the native glycine-specific backbone geometry (**Fig 6D**). This structural distortion, occurring at a residue immediately adjacent to the D166 inter-subunit interaction network disrupted directly by D166N, supports that the T1 inter-subunit contacts identified here represent a structurally vulnerable and functionally significant node under selection in colorectal adenocarcinoma (**Fig 6E**). This is corroborated by elastic network modelling of the WT tetramer, which places both G128 and D166/R167 within 1-3 residues of a predicted dynamic hinge site in the T1 domain (**Fig S2**).

### Transcriptomic Analysis of KCNA5 Mutant Patients Reveals Immune Cell and Proliferative Pathway Changes

To uncover what KCNA5 mutations actually do to tumour biology, we compared transcriptomic (RNA-seq) profiles of KCNA5-mutant and wild-type (WT) COADREAD tumours including MSI score and tumour mutation burden as covariates. Pooling all missense KCNA5 mutations for which we had transcriptomic data regardless of structural domain (n=26) against WT (n=504) identified 179 differentially expressed genes with padj<0.05 (**Fig 7A**). The most significantly altered genes include the water channel aquaporin AQP5 (log2FC=−4.08) and the chloride channel GABRA3 (log2FC=−4.61) both of which were strongly downregulated, and the SLC4A10 pH regulating sodium bicarbonate exchanger which was strongly upregulated.

**Figure 7:**
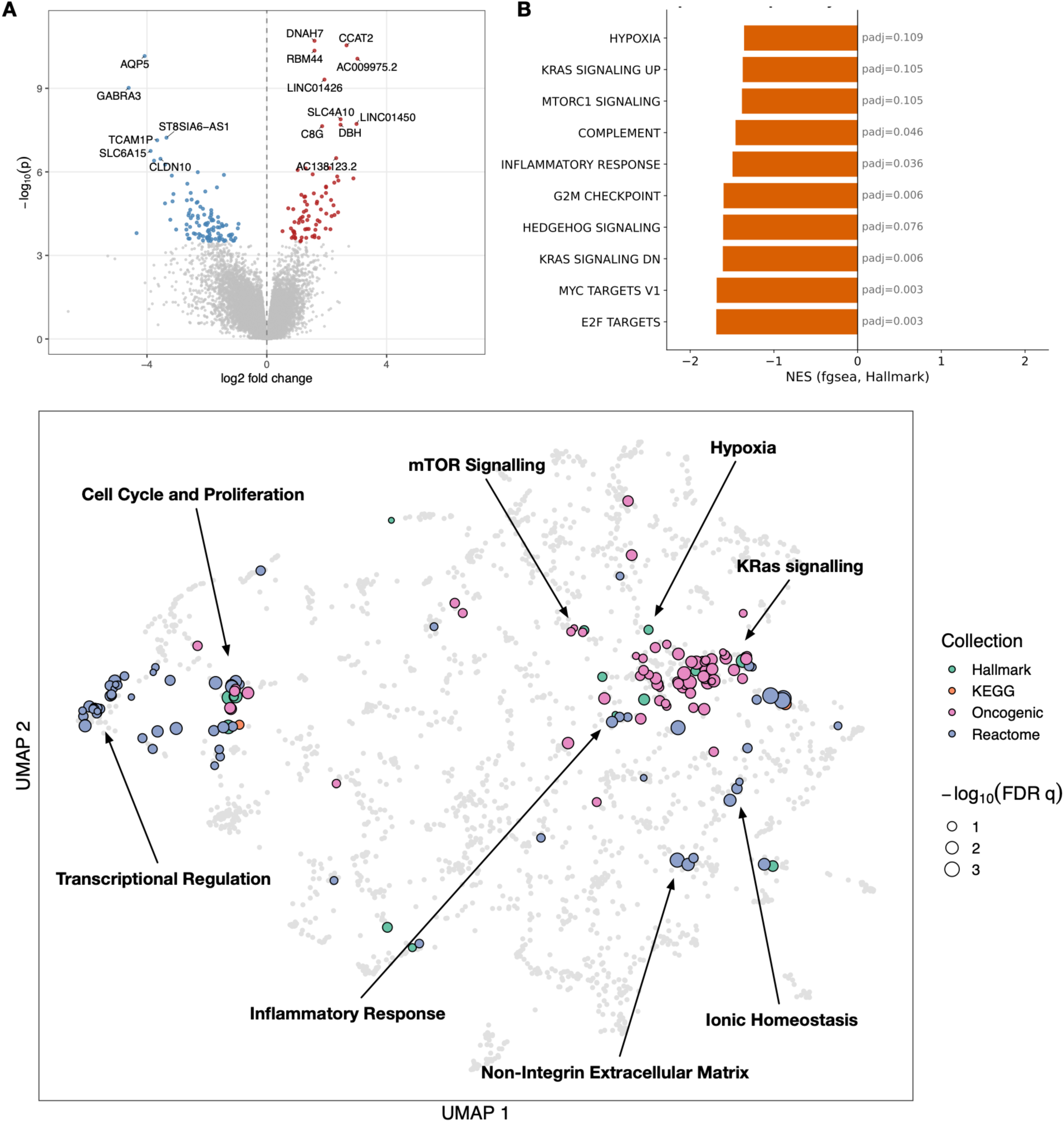
Transcriptomics analysis of KCNA5 mutant vs WT tumors highlights pathways impacted by KCNA5 signalling. A) Volcano plot of gene expression changes in WT vs KCNA5 mutant tumors. B) Significantly differentially expressed pathways as calculated by fGSEA on the Hallmarks pathway set. C) Pathway maps of all significantly (FDR q <= 0.25) differently expressed pathways in 4 gene sets between WT and KCNA5 mutant tumors. Clusters are labelled with their biological activity.

Gene set enrichment analysis (fgsea-Subramanian et al., 2005) against four independent pathway collections - Hallmark, KEGG, Reactome, and the oncogenic signatures set revealed an interesting pattern: of 56 pathways significant at padj<0.05 across all four collections combined (6 Hallmark, 1 KEGG, 21 Reactome, 28 oncogenic), all of them showed a negative enrichment in the mutated patients – indicating that the mutation is correlated with downregulation of these pathways (**Fig 7A**, **Fig S3**). Given that these collections are largely non-overlapping and were tested independently, this directional consistency is likely relevant to mutational effects. Within this negative-NES set, three biologically distinct themes emerged:

1. Reduced proliferation/cell-cycle activity. The strongest and most statistically robust signal in the dataset: Hallmark E2F_TARGETS (NES=−1.69, padj=0.003), MYC_TARGETS_V1 (NES=−1.68, padj=0.003), and G2M_CHECKPOINT (NES=−1.60, padj=0.006), corroborated by approximately ten independent Reactome terms spanning the cell-cycle machinery (DNA_REPLICATION, CELL_CYCLE_CHECKPOINTS, S_PHASE, M_PHASE, MITOTIC_SPINDLE_CHECKPOINT, SEPARATION_OF_SISTER_CHROMATIDS, CHROMOSOME_MAINTENANCE).
2. Reduced KRAS/growth-factor signalling output. Hallmark KRAS_SIGNALING_DN (NES=−1.61, padj=0.006) was reinforced by eight independent KRAS-activity oncogenic signatures (e.g. KRAS.KIDNEY_UP, KRAS.50_UP, KRAS.600_UP), all representing genes normally induced by active KRAS signalling, all showing reduced expression in KCNA5-mutant tumours - consistent with a dampened RAS/MAPK output.
3. Reduced inflammatory/immune pathway activity. Hallmark INFLAMMATORY_RESPONSE (NES=−1.49, padj=0.036) and COMPLEMENT (NES=−1.46, padj=0.046), together with Reactome SIGNALING_BY_INTERLEUKINS, were all downregulated.

We also identified a secondary, less mechanistically resolved signal involved GPCR/receptor-ligand binding, replicated independently in both Reactome and KEGG. Studying the immune pathway signature in more detail, we calculated the immune cell content of the mutant and WT patient samples using xCell[54], and found no difference in immune cell infiltration within the tumor between WT and mutant patients (**Fig S4**) – suggesting that the signature identified is likely a decrease in the immune signalling activity of the tumor itself, rather than a change in the stromal cell population itself.

We also compared transcriptomic signatures from mutated tumors to those from patients with WT KCNA5 in the highest and lowest quartiles of KCNA5 expression to study whether mutations show a similar transcriptomic profile. We find no strong correlation between gene level expression patterns (Spearman ρ= −0.02) between mutation and either over or underexpression patients (**Fig S5**). At the pathway level, we tested concordance across all four gene set collections used elsewhere in this study (Hallmark, KEGG, Reactome, and the oncogenic signatures collection), restricted to the 56 pathways significant in the AllMutant-vs-WT contrast (**Fig S6**). The large majority - 42/56 (75%) - showed the same change in direction between the two contrasts, consistent with expression-dose-dependent effects for pathways such as KRAS signalling, inflammatory response, and complement activity. The 14 discordant pathways, however, were not distributed evenly across biological themes: 13 of 14 were cell-cycle or proliferation-related. In every one of these cases, WT tumours with low KCNA5 expression were markedly more proliferative than KCNA5-high WT tumours, the opposite of the reduced-proliferation signature seen in KCNA5-mutant tumours. This indicates that the divergence between mutation and low expression is not a general phenomenon but is concentrated specifically on cell-cycle control, and that the mutant protein’s effect on proliferation is not attributable to reduced expression alone - more consistent with a specific structural consequence of the mutant channel itself than with simple loss of channel expression.

This divergence is consistent with established biology linking potassium channel activity to cell-cycle control. Membrane hyperpolarization is required for progression through the G1/S checkpoint, and Kv channels are a principal driver of this hyperpolarization [17,55] KCNA5’s role within this framework is not fully understood, but some evidence suggests that rather than promoting proliferation, KCNA5 silencing specifically favours proliferation and stress survival in several malignancies, for example Ewing sarcoma, where silencing of KCNA5 supports proliferation and restoring channel function pharmacologically suppresses it[13] and a related study where Polycomb-repression demonstrates that KCNA5 suppression specifically promotes cancer cell survival under stress conditions[56].

### Mutations in the T1 and S1-S4 domains overlap but have distinct transcriptional effects

To determine whether mutations in different domains of the protein converge on a shared transcriptional programme or induce unique effects, we separated our cohorts into patients with mutaions in different domains of KCNA5. T1 domains and the S1-S4 domain contained 10 patients with missense mutations in KCNA5, whilst the pore/selectivity filter region only contained 1 patients with both a mutation in KCNA5 and transcriptional data. We focussed on T1 and S1-S4 mutations, and we compared their effects at the gene and pathway levels, both directly (T1-mutant vs S1-S4-mutant) and indirectly (each domain vs WT). At the gene level, differential expression overlapped minimally: of the genes significant at padj<0.05 in either domain relative to WT, 34 were T1-specific, 29 were S1-S4-specific, and only one (GABRA3) was shared between domains (**Fig 8A**).

**Figure 8:**
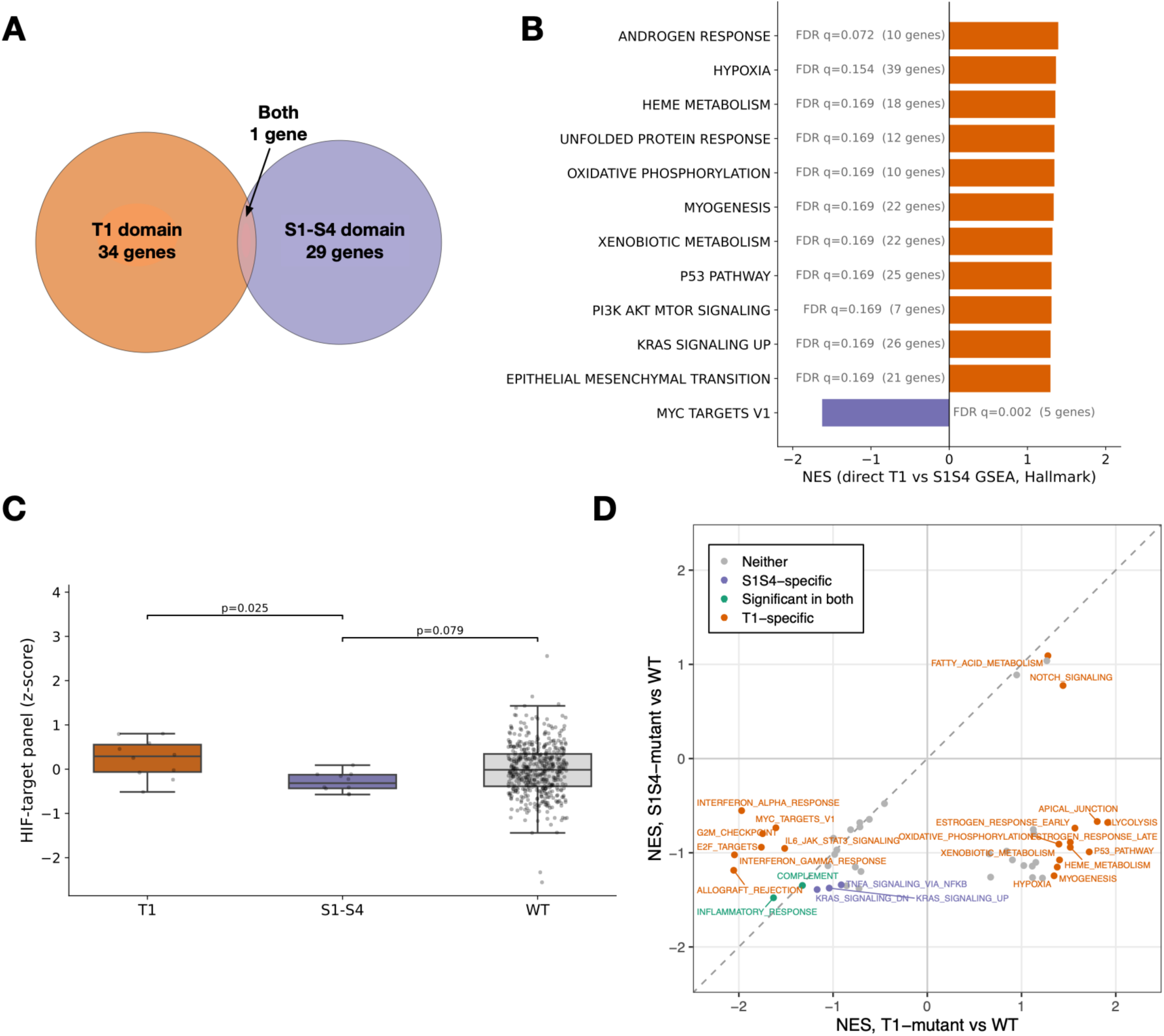
Transcriptomic similarities and differences between different types of KCNA5 mutants. A) Venn diagram of differentially expressed genes between T1 vs WT and S1-S4 domain vs WT. B) Differentially expressed pathways in the Hallmarks set as calculated by fGSEA between T1 and S1-S4 mutant tumors. C) Difference in hypoxia score between T1 and S-S4 mutant tumors. D) Pathway comparison for all pathways in 4 gene sets between T1 and S1-S4 mutant tumors.

A pathway analysis performed as a direct contrast between T1- and S1-S4-mutant tumours however identified 12 Hallmark pathways distinguishing the two domains at FDR q<0.25 (**Fig 8B**). Eleven of these were higher in T1, while genes relating to Myc targets were the single pathway in the hallmarks set higher in S1-S4. One pathway of interest was the Hypoxia pathway – which KCNA5 has been demonstrated to play a role in. To cross-validate the hypoxia signal independently of GSEA’s ranking method, we scored each tumour for a previously curated HIF-target gene panel. T1-mutant tumours scored higher than S1-S4-mutant tumours (p=0.025) and trended higher than WT (**Fig 8C**), consistent with the GSEA result. Together, these results indicate that T1 and S1-S4 mutations perturb a largely shared transcriptional axis, but with T1 producing a substantially broader and stronger signature consistent with T1’s role in the cooperative, tetrameric assembly interface versus S1-S4’s more localised function in voltage sensing.

Finally – to compare the transcriptional effects of each mutation we performed a pathway analysis for each mutation group against WT patients, and compared pathway enrichments. Pathway level changes were broadly correlated between the two mutational categories (Hallmark NES, Spearman ρ=0.276, p=0.052)(**Fig 8D**), and the majority of pathways with small effect sizes fell on the same side of zero in both mutation groups, but some pathways showed opposite trends. KRAS and TNF alpha pathways appeared to be more specific to S1-S4 domain mutations, whilst a broader set of pathways was specific to T1 mutations, including interferon signalling, proliferation-related and several metabolic pathways (19 T1-specific vs 3 S1-S4-specific Hallmark pathways at FDR q<0.25). Only INFLAMMATORY_RESPONSE and COMPLEMENT were significant in both mutation groups. T1-specific pathways also tended to have larger effect sizes than the corresponding, non-significant S1-S4 values, indicating that T1 and S1-S4 mutations act on a largely overlapping set of pathways, but that T1 mutations produce a stronger and more extensive transcriptional response.

This asymmetry is consistent with the two domains’ distinct structural roles. S1-S4 forms the voltage-sensing domain, and its charged gating-charge residues are specifically enriched for charge-disrupting mutations in this cohort, pointing to a precise biophysical mechanism - impaired voltage sensing - that need not perturb bulk transcription to have a phenotypic effect. T1, by contrast, forms the N-terminal tetramerisation domain that governs cooperative assembly of all four channel subunits; disrupting this interface plausibly compromises proper channel folding and assembly, consistent with UNFOLDED_PROTEIN_RESPONSE emerging as one of the pathways most specific to T1-mutant tumours. Overall this points to mutations in different domains of the protein having broadly overlapping but partially distinct effects related to their effects on the protein.

## Conclusions

Together, these results support a model in which KCNA5 is a positively selected driver gene in colorectal adenocarcinoma. Mutations are non-randomly distributed across three structurally and functionally distinct domains, each domain’s mutations have a plausible, domain-appropriate biophysical consequence (loss of gating-charge function and a putative gating-pore leak in S1–S4; destabilisation of inter-subunit contacts and interface dynamics in T1; a sterically disruptive but filter-sparing biophysical signature in the pore domain), and these structural predictions are mirrored by a coherent, directionally consistent transcriptional phenotype (reduced proliferative, KRAS, and inflammatory signalling) that is not reproduced by reduced KCNA5 expression alone.

The pore-domain finding is the most unexpected: cancer-selected mutations avoid the selectivity filter and the previously characterised classes of gating-altering mutation, instead occupying what appears to be a distinct biophysical niche, suggesting that whatever selective advantage these mutations confer, it is unlikely to operate through simple channel blockade or loss of conduction. A degree of caution is warranted in interpreting these findings: the structural and energetic predictions, while internally consistent and in several cases corroborated against experimentally characterised homologous positions in KCNA2, remain computational rather than electrophysiologically validated, and the transcriptomic analysis - correlative by nature cannot establish causality or fully separate domain-specific mutational effects from unmeasured confounders. Direct electrophysiological characterisation of representative mutations from each domain, particularly the predicted gating-pore candidates (R403H, R409H) and the pore-domain mutations whose biophysical class has no existing functional correlate, would be necessary to confirm the mechanisms proposed here. Nonetheless, this work demonstrates that combining selection analysis with structure-based biophysical modelling and transcriptomics offers a tractable route to generating testable, residue-level functional hypotheses for cancer-associated ion channel mutations at a scale that direct electrophysiology cannot match, and identifies KCNA5 as a clear candidate for such follow-up study in colorectal cancer.

## Methods

### Data acquisition

COSMIC data were downloaded from cancer.sanger.ac.uk (version 103) [57]. We subset mutations into those only found in the colorectal adenocarcinoma, with the filter: “large intestine”, “include all”, “carcinoma”, “adenocarcinoma”.

RNA seq data from the GDC TCGA Colon Cancer (COAD) dataset was downloaded from the UCSC Xenabrowser [58].

Protein structure of KCNA5 was obtained from the SWISS-MODEL predicted structure at UNIPROT, with identifier P22460_119-527:2r9r.3.B (https://swissmodel.expasy.org/repository/uniprot/P22460?range=119-527Ctemplate=2r9r.3.B).

Protein sequences of the KCNA family were obtained from UNIPROT [59].

### dN/dS Analysis

dN/dS for individual genes was calculated using the dndscv library [60] applied to the cosmic dataset across all genes.

### NMC

Mutational clustering was calculated using the NMC method from the R library IPAC [61] applied to the sequence alone.

### Sequence Alignment

Multiple sequence alignment was done using the Clustal Omega Program [62].

### Generation of Mutant Structures

Mutations were made using the mutate_model protocol in the modeller tool [63] as described in [64], and using the mutate_model.py script available on the modeller website. The structures (both wild type and mutants) were energy minimised using the FoldX command ‘repairpdb’, performed ten times in sequence with the scripts from [65].

### Binding Energy Calculation

Binding energy was calculated using the method described in [65].

### Change in Activation Potential Data Collection

The changes in activation potential were taken from figure 4A in [52] and extracted with PlotDigitizer [66].

### MD Simulation

The simulation systems were generated in CHARMM-GUI bilayer builder from the online server [67]. Lipid bilayers with 100% palmitoyl-oleoylphosphatidylcholine (POPC), containing 200 POPC in the upper leaflets and 215 POPC in the lower leaflets, were generated and the systems were solvated with TIP3 model and completed with 0.8 M K+ and Cl−, reaching an electrically neutral system. The K+ ions in the selectivity filter were preserved in the structure. The wild type system contained 1023 K+ and 1023 Cl−, while the mutant systems contained approximately equal numbers of K+ and Cl− ions, slightly altered to maintain the neutrality. We used the CHARMM36 forcefield[68]

Molecular dynamics was performed using GROMACS, version 2021.5 [69]. The systems were first subjected to 5000-step minimization with the steepest descent algorithm integrator. The systems were further equilibrated in six successive steps with an iteratively decreasing force constant for the positional restraints for a total equilibration time of 1.875 ns and starting from randomly assigned velocities. The first initial two equilibration steps were run as NVT ensemble, before switching to the NPT ensemble for the remainder of the equilibration as well as production runs. Temperature coupling at 303.15 K was achieved using the v-rescale thermostat, while pressure coupling was achieved using the c-rescale barostat.

For the production runs, without positional restraints, random velocities based on the Boltzmann distribution were assigned to both systems and unbiased simulations were run with a time step of 2 fs. Hydrogen bonds were constrained using Linear constraints solver (LINCS) and long-range electrostatic interactions were computed using Particle-Mesh Ewald (PME) with 1.2 nm cutoff.

### Differential Expression Analyses with DESeq2

Differential gene expression was performed using DESeq2 (v1.40) [70] in R on the GDC TCGA Colon Cancer (COAD) RNA-seq raw counts data. Three contrasts were analysed: DESeq2 between the tumour and normal samples were computed to study the change in expression of KCNA5, and two extra analyses (KCNA5 T1 domain mutant (residues 128-216) vs confirmed wild-type tumours, and non-T1 KCNA5 mutant vs confirmed wild-type tumours) were calculated for the gsea analyses. Size factors were estimated using the median-of-ratios method and dispersion was estimated using the default DESeq2 workflow. Results were extracted using a Wald test with Benjamini-Hochberg multiple testing correction, with a significance threshold of padj < 0.05.

### Gene Set Enrichment Analysis (GSEA)

GSEA was performed using the fgsea package (v1.26) [71] in R with 10,000 permutations and exact p-value calculation (eps = 0). Genes were ranked by the metric:

rank = sign(log2FoldChange) × −log10(adjusted p-value)

capturing both the direction and statistical confidence of differential expression. P-values equal to zero were replaced with the smallest representable double-precision floating point number to avoid infinite rank values. The Hallmark collection from MSigDB [72] was used for analysis. Gene set collections were accessed using the msigdbr package (v7.5). Pathways with Benjamini-Hochberg adjusted p-value < 0.05 were considered significant.

## Acknowledgements

The authors acknowledge the use of the UCL Myriad HPC platform, and the support of the UCL Centre for Advanced Research Computing, in the completion of this work.

