## Supplementary Information for "Positive selection and structural disruption of KCNA5 by mutation in colorectal cancer"

**Supplementary Information for Xu et al. Positive selection and structural disruption of KCNA5 by mutation in colorectal cancer**

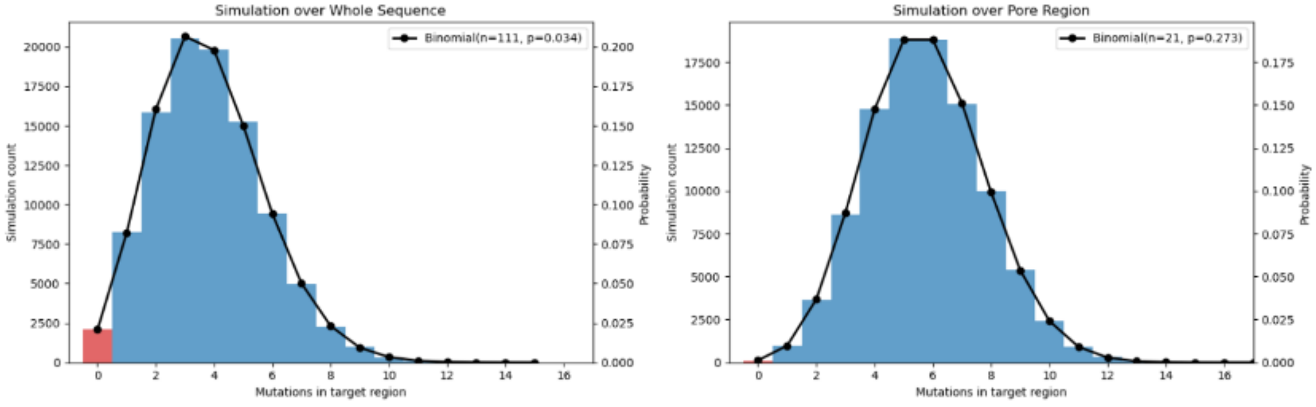


Supplementary Figure 1: Results of Monte-Carlo simulations of mutations to the whole protein (left) and pore region (right), with zero values coloured red.

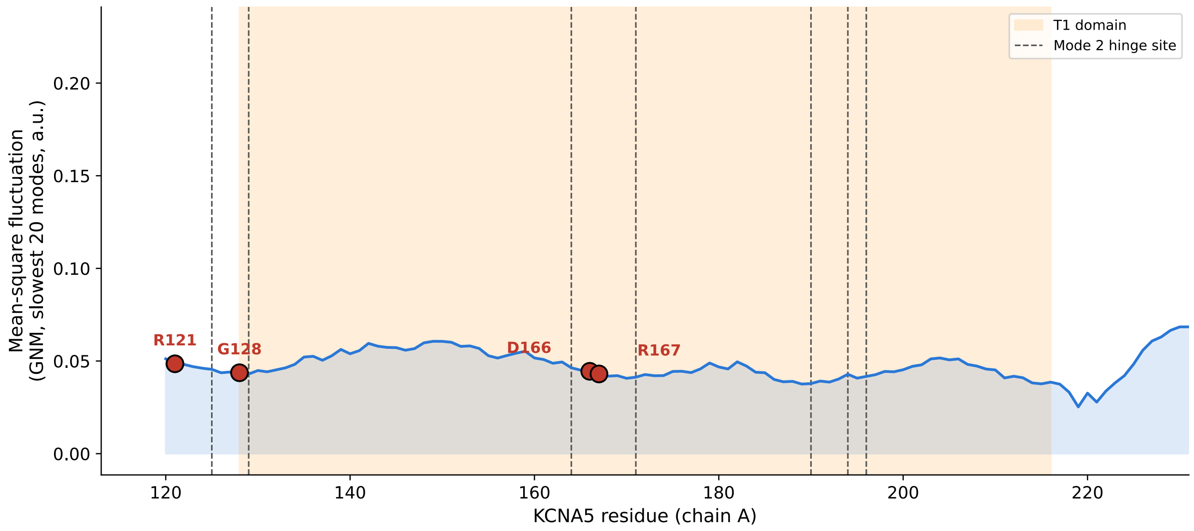


Supplementary Figure 2: Elastic Network mode 2 hinge sides identified along the protein sequence, with residues that are recurrently mutated highlighted.

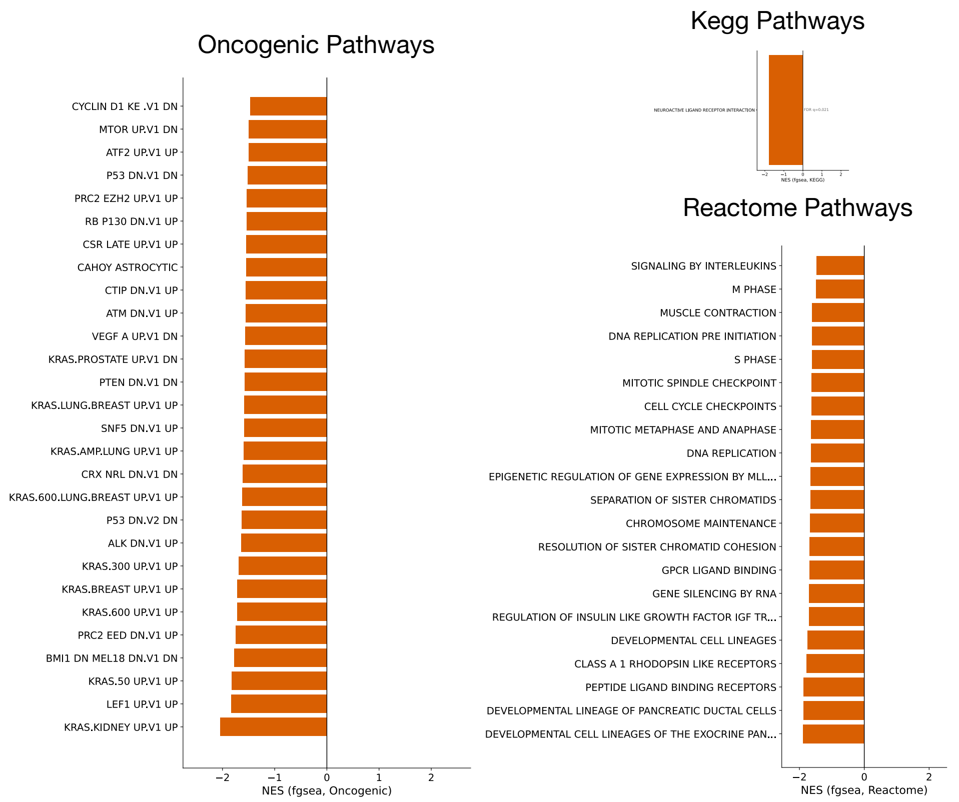


Supplementary Figure 3: Pathways differentially expressed between KCNA5 mutant vs WT tumours.

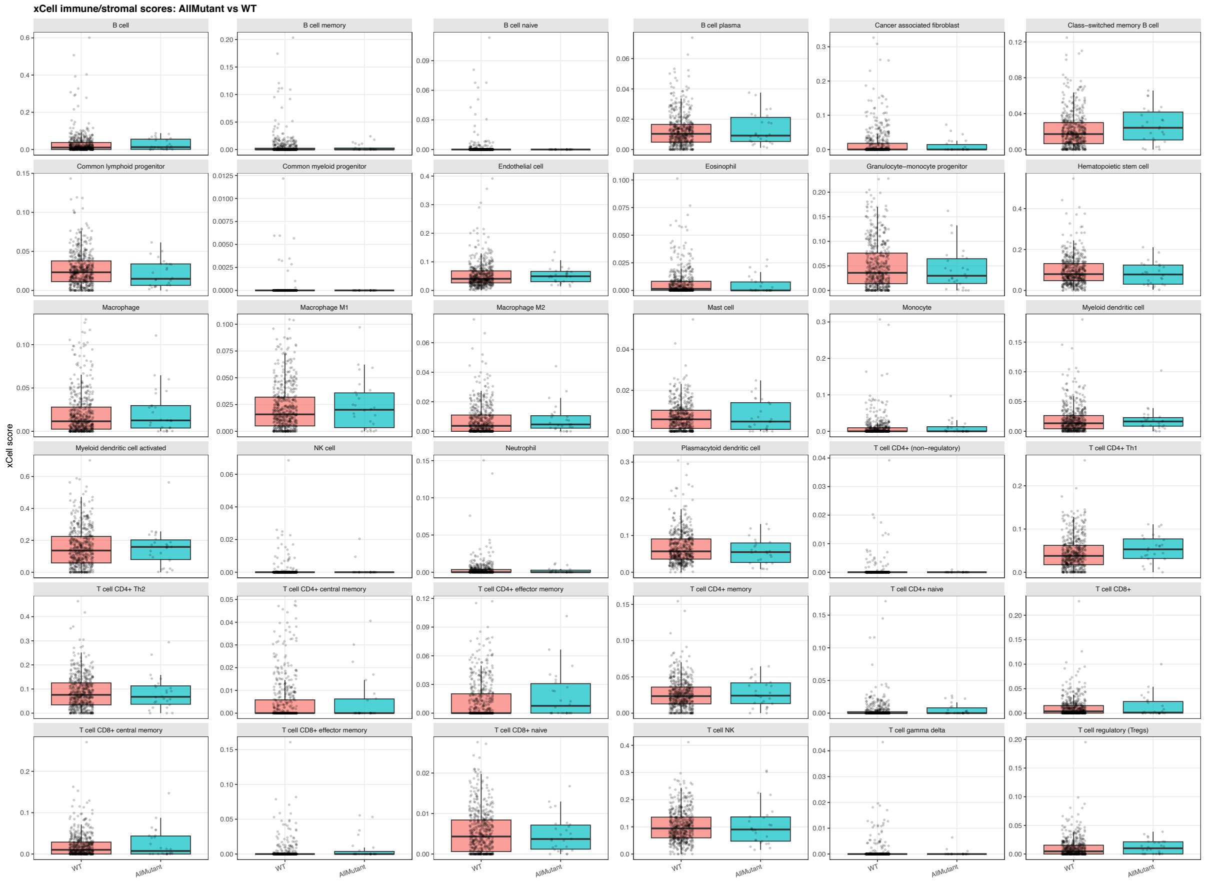


Supplementary Figure 4: xCell deconvolution for WT and KCNA5 mutant tumours

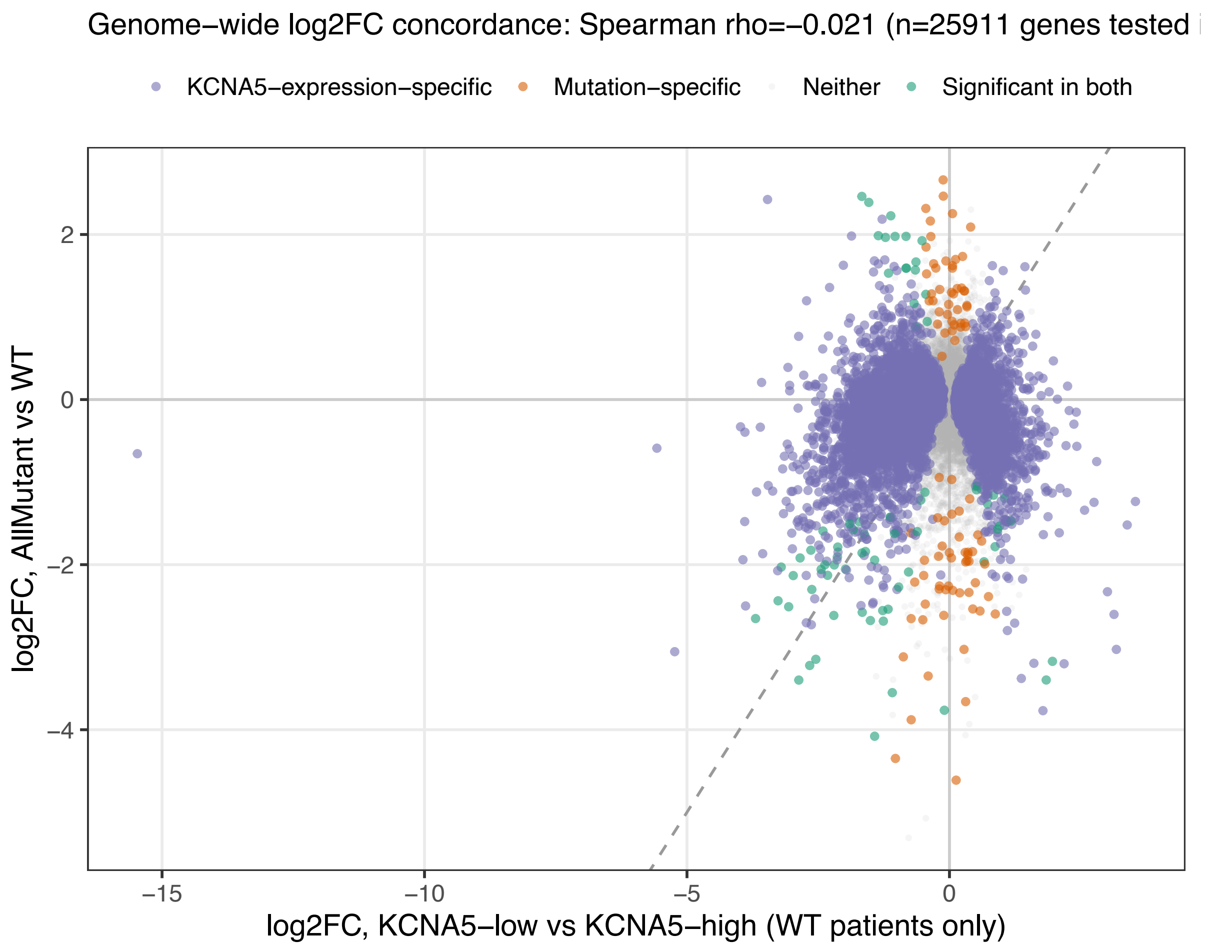


Supplementary Figure 5: Concordance analysis of gene expression between KCNA5 mutant vs WT tumours, and KCNA5 low vs KCNA5 high expression tumours.

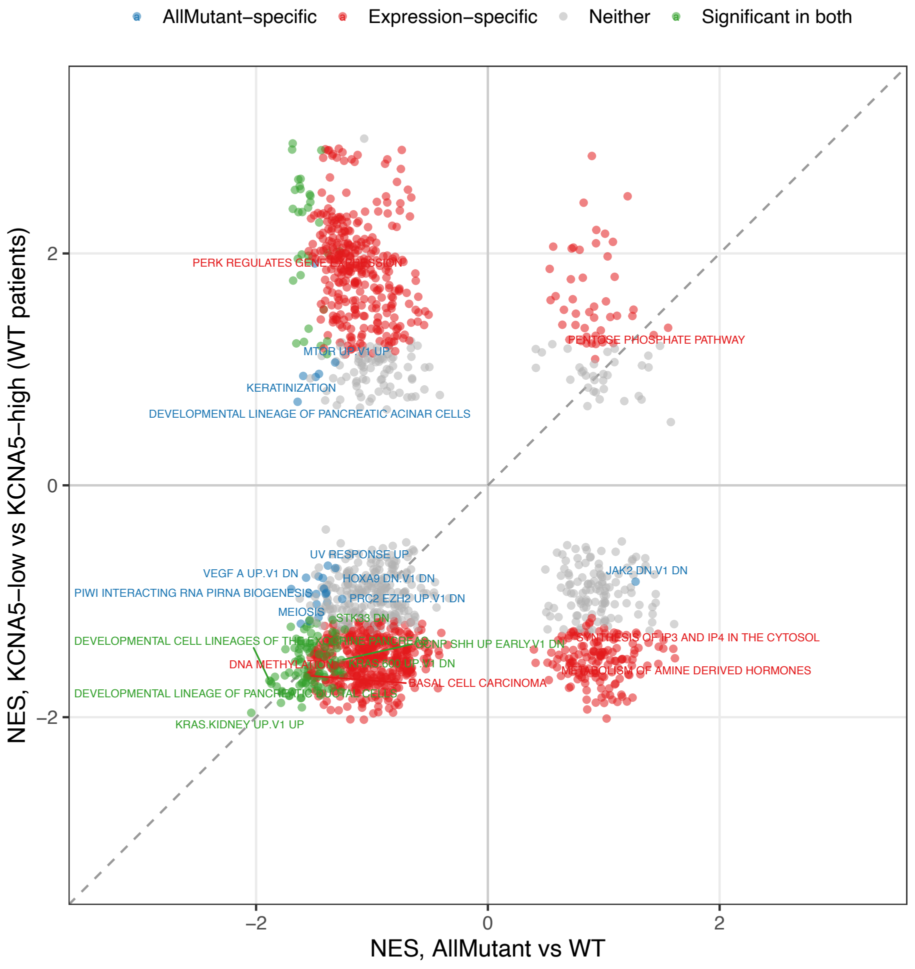


Supplementary Figure 6: Concordance analysis of pathway enrichment between KCNA5 mutant vs WT tumours, and KCNA5 low vs KCNA5 high expression tumours. Shown are all pathways with FDR q <= 0.25 in either analysis.
